# Population dynamics of *Diatraea* spp. Under different climate offer conditions in Colombia

**DOI:** 10.1101/2020.02.20.957977

**Authors:** Julián Andrés Valencia Arbeláez, Alberto Soto Giraldo, Gabriel Jaime Castaño Villa, Luis Fernando Vallejo Espinosa, Melba Ruth Salazar Gutiérrez, Germán Andrés Vargas Orozco

## Abstract

Seasonal temperature and precipitation patterns on a global scale are the main factors to identify the sharing of organisms. Accordingly, insects and plants come to adapt to combinations of various factors through natural selection, although periodic outbreaks in insect populations occur especially in areas where they have not been previously reported, is a phenomenon that is considered as a consequence of global warming. In the study, we sought estimate the distribution of the sugarcane stem borers, *Diatraea* spp., under different climate scenarios. Weekly collections were carried out in four sugarcane field plots in four different towns from the Colombian department of Caldas during a consecutive year, and also from sugarcane plots from the Cauca river valley between 2010 and 2017. The influence of climatic variables on the climate in different agro-ecological zones of sugarcane crops (*Saccharum* sp.) was defined by using climatic data (maximum, minimum and daily temperatures; accumulated precipitation) on a daily scale between 2010 and 2017. MODESTR® was used to generate the distribution maps to estimate probability distributions subject to restrictions given by the environmental information. *Diatraea* spp. is strongly influenced by the effects of climate change, considerably reducing its population niches as well as the number of individuals. The estimate of an optimal niche for *Diatraea* spp. includes temperatures between 20°C and 23°C, accumulated annual rainfall between 1200 and 1500 mm, months with dry conditions, whose precipitation is below 50 mm, slopes below 0.05, crop heterogeneity with an index of 0.2 and primary production values of 1.0.

**Summary statement:** *Diatraea* spp. is susceptible to temperature variations due to climate change, it is presumed that its adaptability could benefit *Diatraea* spp. in establishing itself in new areas.

## Introduction

The Earth’s temperature increased between 0. 7°C from 1900, 1. 3°C from 1950 and 1. 8°C over the past 35 years. The last two decades have been among the warmest since temperatures began to be recorded. Since 1981, annual tons of agricultural crops have been lost due to global warming (Peng *et al*., 2004; Lobell and Field, 2007), including the sugarcane crop; although in some cases, where technification is evident, these losses were compensated for by higher yields achieved through genetic improvements in crops and other agrotechnological advances (Foley *et al*., 2011).

It would that, beyond what has been noted over the past 50 years, high seasonal temperatures may become even more widespread in various parts of Mesoamerica and South America during the remainder of this century (Battisti and Naylor, 2009). It is estimated that temperatures could rise by 0. 4°C to 1. 8°C by 2020, and this increase would be even more pronounced in tropical areas. High temperatures (especially when increases exceed 3°C) affect significantly agricultural productivity, producer incomes and food security. A number of crops that represent essential food sources for large food-insecure populations will have a serious impact on their yields, although it appears that the scenarios are more uncertain for some crops than others (Nelson *et al*., 2011; Peng *et al*., 2004).

In agricultural systems, there have been increases or decreases in incidence of pests associated with extreme events of climate change, such as prolonged droughts, hurricanes, heavy and out of season rains, among others. Of course, these are often not noticeable, because the disasters caused by such events to crops do not allow us to appreciate the changes in the manifestations of pests. However, these contribute to increased losses, forcing farmers to spend too much on pesticides that usually fail to solve the problem. (Estay, 2009; Vázquez, 2011).

On a global scale, seasonal patterns of temperature and precipitation are known to be the main factors in determining the distribution of organisms (Birch, 1948; Marco, 2001). Insects and plants come to adapt to combinations of these factors through natural selection, although insects with periodic outbreaks occur especially in areas that are physically severe or stressed, a phenomenon that is considered a consequence of global warming. (Vázquez, 2011).

Due to the insect populations respond to changes in local climate or levels of sun’s light, humidity, precipitation and temperature, it has been suggested that changes in these populations may serve as an indicator of local or global climate change (Marco, 2001). But temperature is the event that affects most due mainly to its important impact on biochemical processes, since arthropods cannot regulate their body temperature, depending for the environmental temperature to obtain heat from exposure to solar radiation (Wagner *et al*., 1984), directly affecting the dynamics of infestation (Marco, 2001; Jaramillo *et al*., 2009, Estay; 2009), and these can be used as ideal bioindicator for an agricultural crop. In addition, the increasingly pronounced climate variability events (El Niño and La Niña) have been a cause of study for the scientific community, being the main topic of the research scenarios, concluding among many discussions that arthropods are living organisms mostly considered as bioindicators of changes in climate behavior, since most insects have a rate of development that is limited mainly by temperature changes, altitudinal differences and their distribution is directly associated with their eating habits (Barbosa y Perecin, 1982; Battisti y Naylor, 2009; Chen *et al*., 2014). These factors are combined to develop agrometeorological indices, which allow the identification of areas of greater and lesser risk of climate variability, while helping with regionalization recommendations for future climate variation events (Muñoz, 2012). Therefore, monitoring studies with a bio-indicator of climate change can play a role with two main elements: First, the data obtained can be used to predict the effects of climate change, and second, monitoring over a long period of time can detect changes in abundance and distribution and establish management strategies that allow the adoption of appropriate control measures, becoming valuable sources of information on the effects of climate change in tropical latitudes, where there is not enough information.

The species belonging to the genus *Diatraea* in sugarcane (*Saccharum* sp.) are those of greatest economic importance in the Americas, since their presence is permanent, and its effects are expressed on either biomass production and/or sucrose content (Lange *et al*., 2004). The damage and losses caused by the insect can be described as destruction of buds in planting material; in seedlings, damage to the bud causing the so-called dead heart; circular perforations in the nodes and internodes that cause the cane to break and allow the entry of other insects or diseases, decrease in the sucrose content due to the inversion process suffered by sugars through the harmful action of the borer and other pathogenic organisms, among others (Vargas *et al*, 2015). However, its importance and the numerous studies on different aspects of biology, integrated insect management and control methods there is a lack of research on aspects of population dynamics and insect ecology, particularly related to the effect of climate change and its dispersion.

In order to understand the effect that climate change will have on the distribution of species of economic importance for sugarcane crops, we seek to generate predictive models on inter- and intra-annual scales at the global, national and local levels, which will allow decisions to be made on management and control practices, thus reducing levels of uncertainty in the face of different climate scenarios.

## Materials and methods

Records from the different lots of each sugar mill in the geographic valley of the Cauca River (which extends from the north of the department of Cauca, across the center of the department of Valle and into the south of the department of Risaralda) and from the department of Caldas, Colombia between the years 2010 and 2017 were used. All the collections followed the same methodology in the field: 120 points (canes) chosen at random during one year, making a zigzagging route within the cane field during 40 minutes per plot, following the orientation according to the linear distribution of the cane (Pérez *et al*. 2011), where samples of eggs, larvae, pupae and adults were taken, from the moment the first true leaves appear in the sugarcane crop until the time of harvest. Collections were manual in each of the sites sampled, in order to obtain the intensity of infestation, recorded as the location of the collection points, so that a representative sampling of the area can be obtained. The global data were obtained from the invasive compendium of CABI species (www.cabi.org),

The climatic variables influence on the insect development in different agro-ecological zones was defined, resorting to the use of the climatic data (maximum, minimum and daily temperature; precipitation) at daily scale between 2010 and 2017 provided by the Colombian Sugarcane Research Center – Cenicaña (Fig 1).

**Fig 1.**
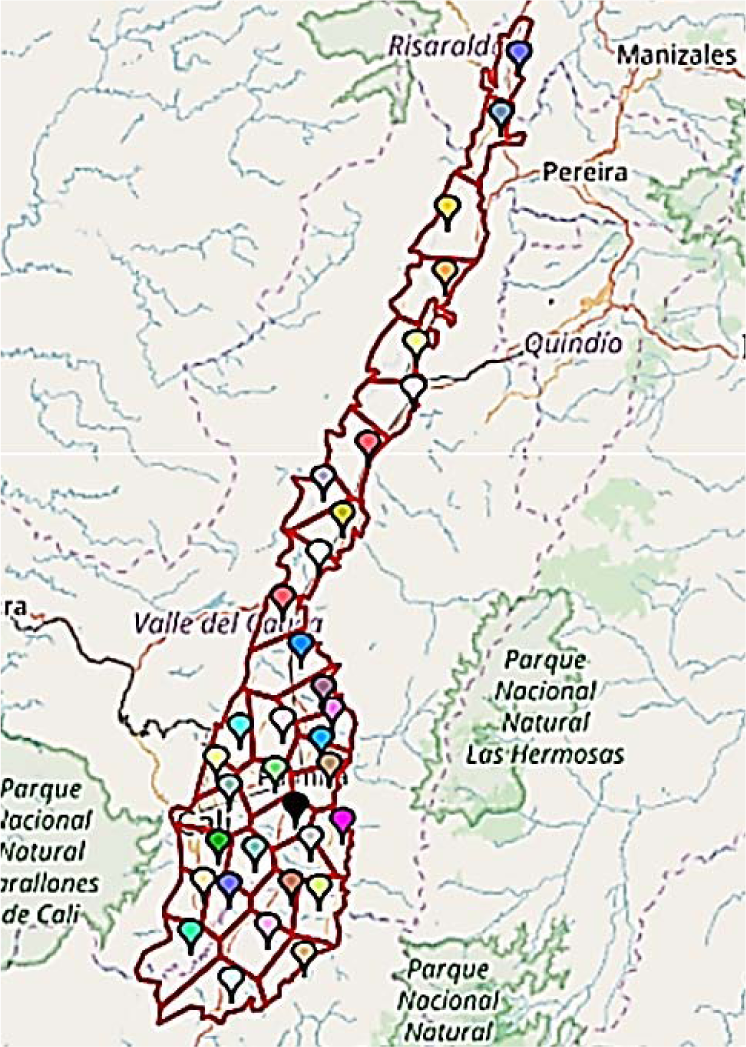
Location of the weather stations in the geographic valley of the Cauca River, Colombia

To generate the missing weather data, Marksim® was used, which provides daily scale weather data files. (http://gismap.ciat.cgiar.org/MarkSimGCM/). At the same time, open-access records available on weather portals were searched for the use of WorldClim and collected the 19 bioclimatic variables derived from monthly rainfall and temperature values (Hijmans *et al*., 2005).

To determine the level of correlation between the variables, the VIF (Variance Inflation Factor) was calculated (Estrada *et al.,* 2013; Pradhan, 2016), providing an index that measures the extent to which the variance (the square of the estimated standard deviation) increases due to the collinearity, allowing the discarding of those climatic variables that can generate noise through redundancy (Equation 1).

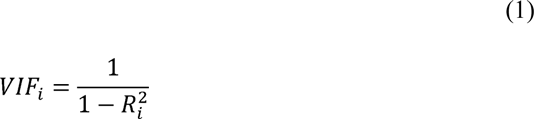

Where 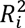 is the determination coefficient of the regression equation

Finally, only 5 of the 19 variables provided by WorldClim plus Standardized Vegetation Index, slope, altitude, aspect and topographic heterogeneity were used (Larkin *et al.,* 2005; Zhang *et al*., 2017), supplied within the ModestR® software (Table 1).

**Table 1.**
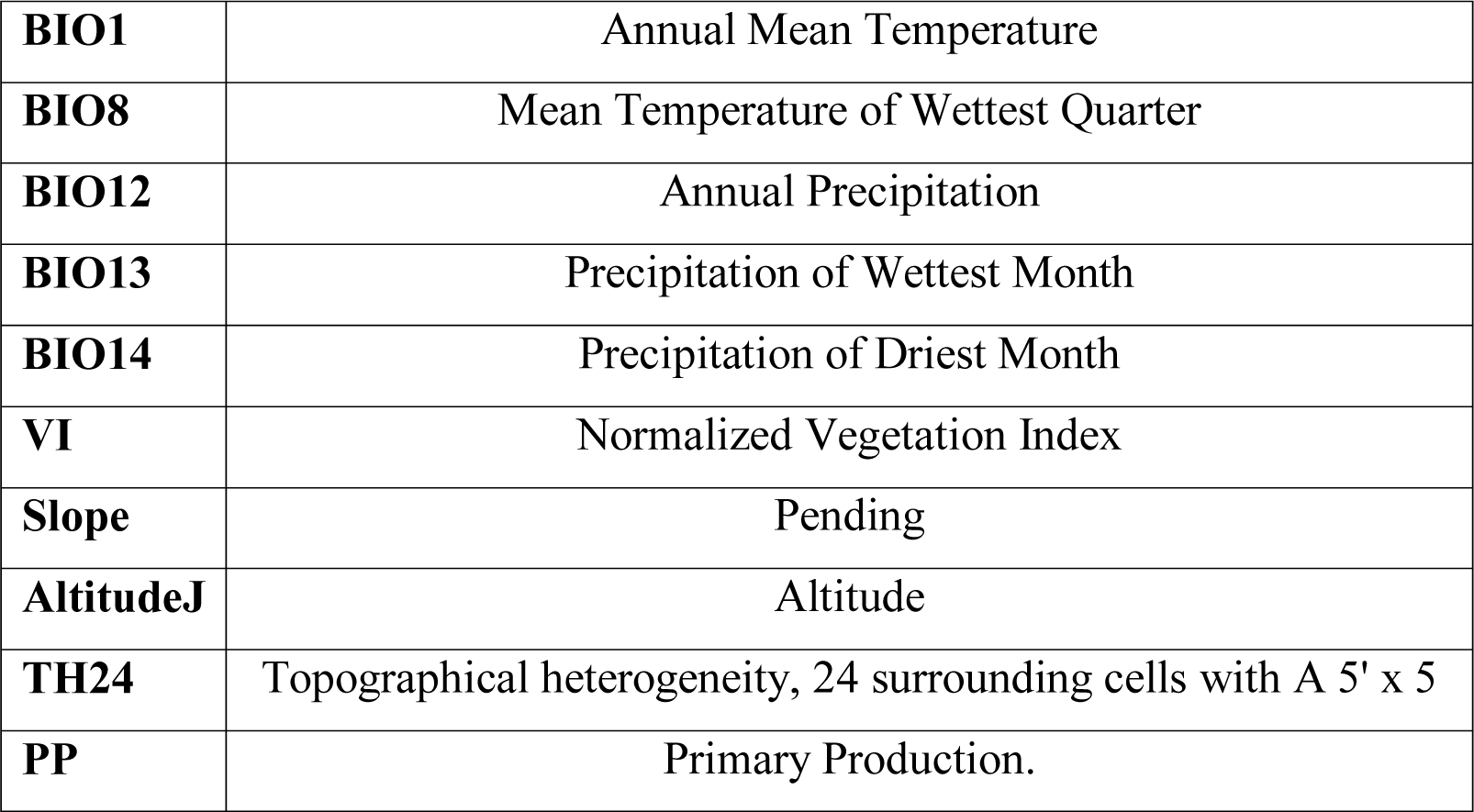
WorldClim Climate variables

|  |  |
| --- | --- |
| **BIO1** | Annual Mean Temperature |
| **BIO8** | Mean Temperature of Wettest Quarter |
| **BIO12** | Annual Precipitation |
| **BIO13** | Precipitation of Wettest Month |
| **BIO14** | Precipitation of Driest Month |
| **VI** | Normalized Vegetation Index |
| **Slope** | Pending |
| **AltitudeJ** | Altitude |
| **TH24** | Topographical heterogeneity, 24 surrounding cells with A 5' x 5 |
| **PP** | Primary Production. |

The climate change model used was BCC_CSM1 from the Beijing Climate Centre; this is a fully coupled global climate and carbon model that includes the interactive vegetation and global carbon cycle, the oceanic component, the terrestrial component and the sea-ice component, which are fully coupled and interact with each other through impulse flows, energy, water and carbon at their interfaces. The information between the atmosphere and the ocean is exchanged once a day simulated. The exchange of atmospheric carbon with the terrestrial biosphere is calculated at each stage of the model (20 min). For the model, representative concentration pathways (CPR) were used. These are four paths of greenhouse gas concentration (non-emission) adopted by the IPCC and are used to model climate by describing four possible future climate scenarios, which are considered possible depending on the amount of greenhouse gases emitted in the coming years. Reference was made to rcp26, rcp45, rcp60, rcp85. (Wu *et al*., 2012).

To generate the distribution maps and to see the effect for each variable in the distribution of *Diatraea* spp., MODESTR® (http://www.ipezes/ModestR) was used; this is a tool that combines statistics, maximum entropy and Bayesian methods, whose purpose is to estimate probability distributions subject to restrictions given by environmental information. MODESTR®, like MAXENT®, (Elith *el al*., 2011, Phillips and Dudik, 2008) it requires only presence data, and uses continuous and categorical variables, it can incorporate interactions between different variables, in linear models and additives without interaction, allows the evaluation of the role of each environmental variable, while over-adjustment can be avoided using regularizations, the output variable is still allowing distinctions between areas (although it allows discrete), and can be used in multiple applications and on all scales. (Edelstein *el al*., 2010).

To calculate the variable effects of richness, niche, and construction we used RWizard®. The packages used were FactorsR to identify the factors that affect species richness; EnvNicheR to estimate the niche of multiple species and the environmental conditions that favor greater species richness; SPEDInstabR to obtain the relative importance of the factors that affect species distribution based on the concept of stability.

## Results

### Environmental layer

The global environmental layer was generated with each of the variables, using information from WorldCLim (Fig. 2) and a local layer with data from weather stations and Marksim® (Fig. 3), to enhance the distribution model. This layer contains the variables BIO1, BIO8, BIO12, BIO13, BIO14, VI, SLope, AltitudeJ, TH24, PP, with 5×5 cells and in current conditions.

**Fig. 2.**
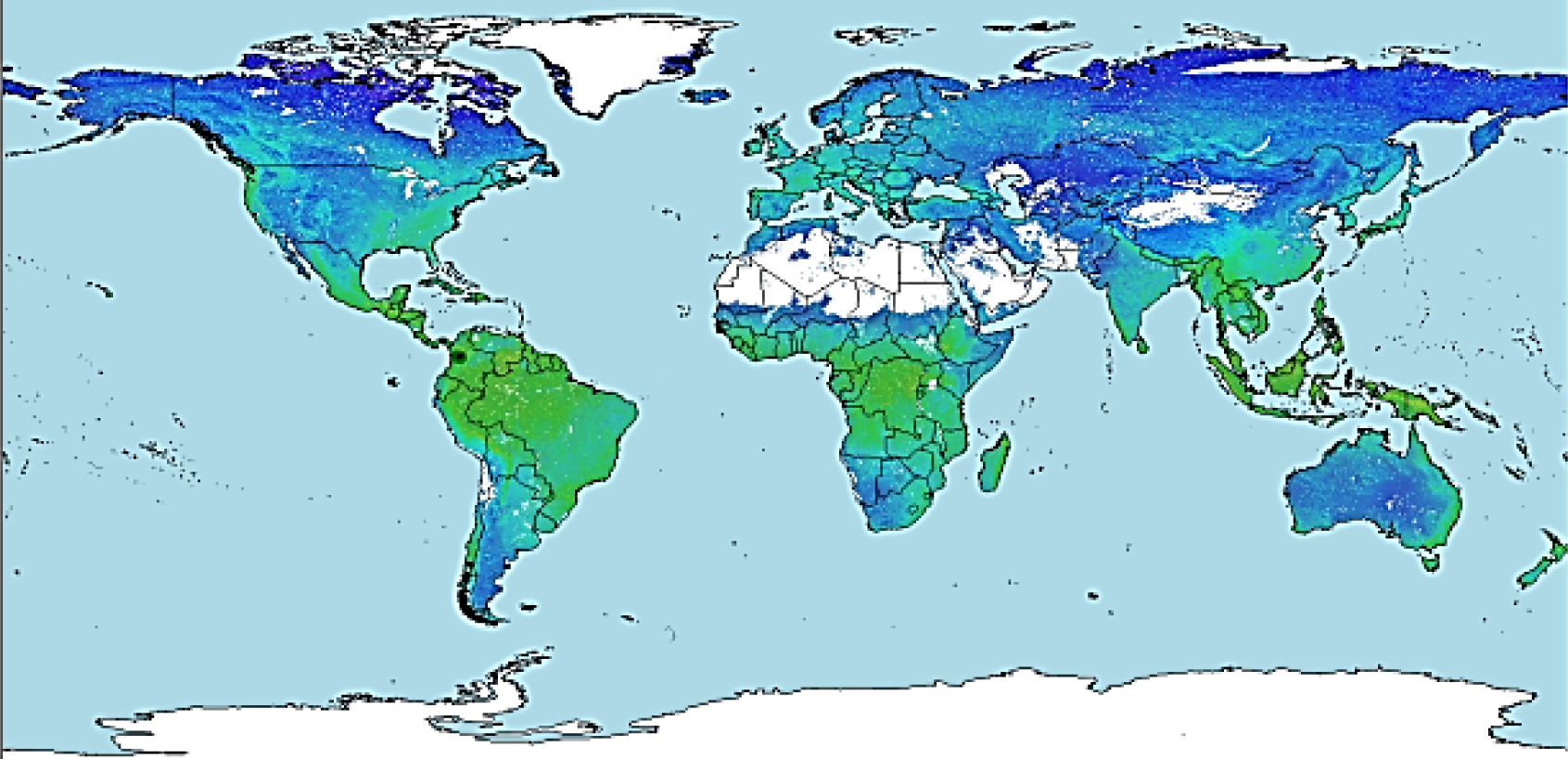
Global scale environmental layer using Worldclim. The colours indicate the mixture of BIO1, BIO8, BIO12, BIO13, BIO14, VI, SLope, AltitudeJ, TH24, PP for each zone in a range of 5’x5’ cell resolution

**Fig. 3.**
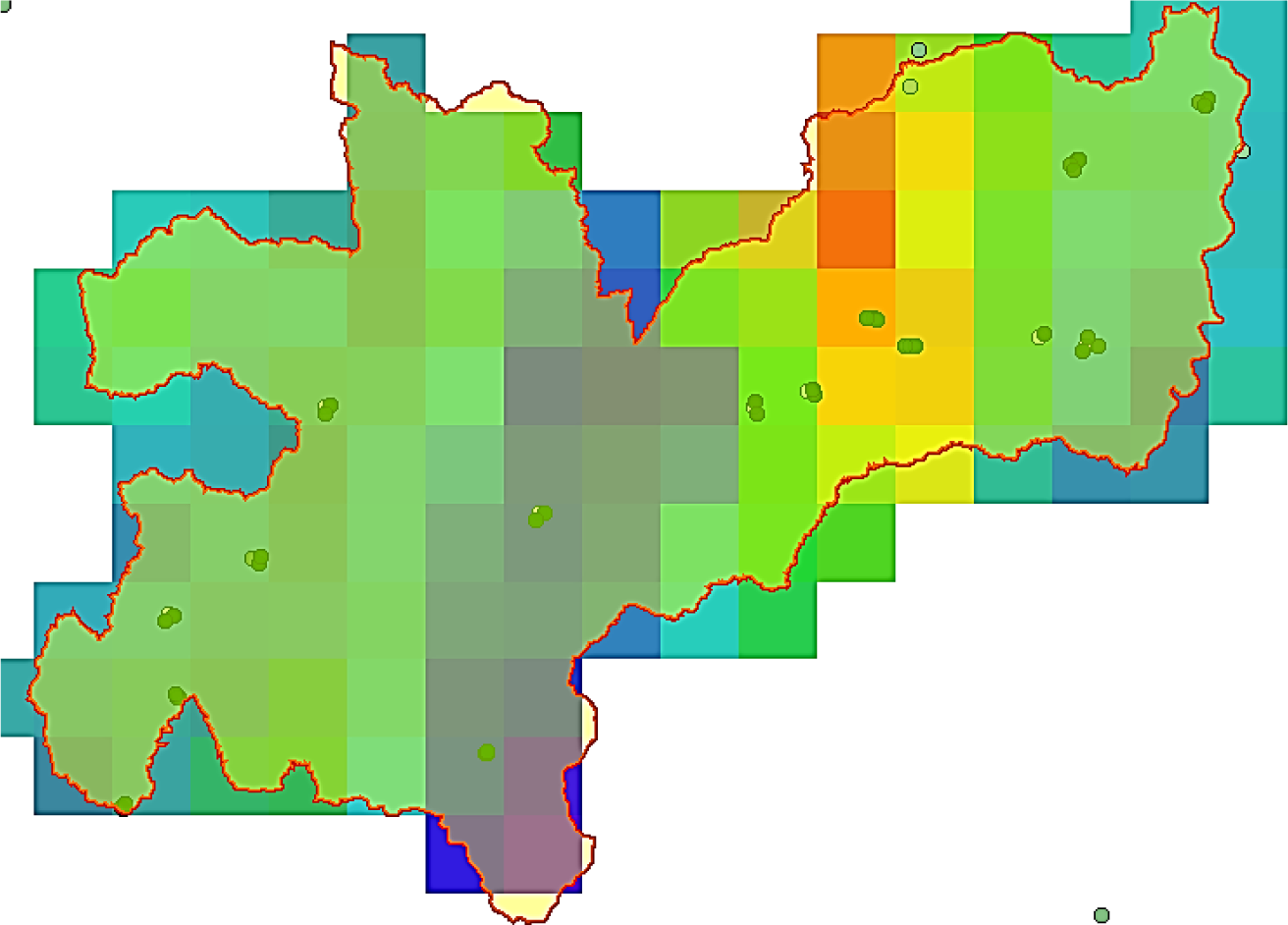
Departamental (Caldas) environmental layer using weather stations and Marksim®. The colours of BIO1, BIO8, BIO12, BIO13, BIO14, VI, SLope, AltitudeJ, TH24, PP for each zone in a range of 5’x5’ cell resolution. The dots indicate records of *Diatraea* spp.

The data obtained were included to generate presence maps for *Diatraea* spp. at global, national and local scales. The local data were records of presence/absence at the sampling points during the research (Fig. 4).

**Fig. 4.**
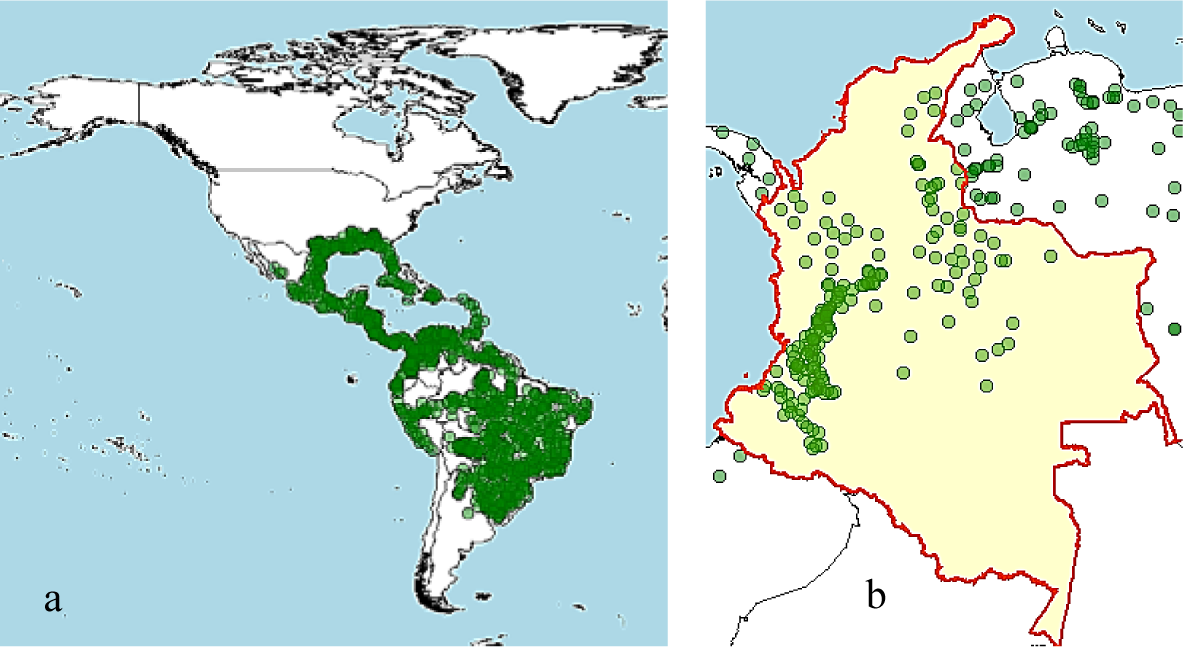
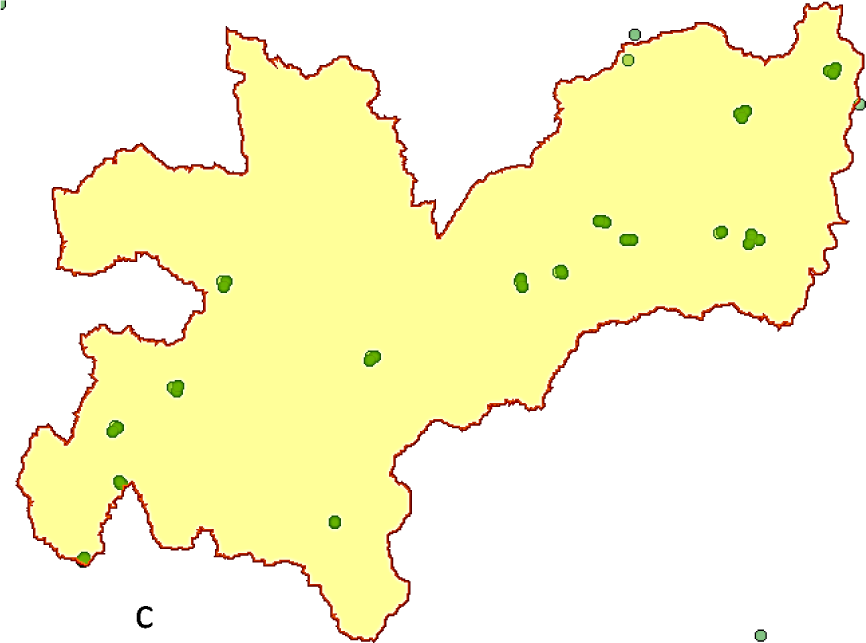
Presence points of *Diatraea* spp. at different scales a) Western hemisphere, b) Colombian and c) Department of Caldas. The dots indicate records

### Variable contribution

The results of the contribution of an environmental variable in the distribution of *Diatraea* spp., show the direct incidence of BIO12, VI, BIO19, PP, SLOPE (slope) and BIO1. Although temperature is the main element for the physiological development of the insect, precipitation is a determining factor in limiting movement and distribution between cultivated plots, as is the availability of food throughout the year and the type of topography. Altitude did not constitute a limiting factor for pest distribution. (Fig. 5).

**Fig. 5.**
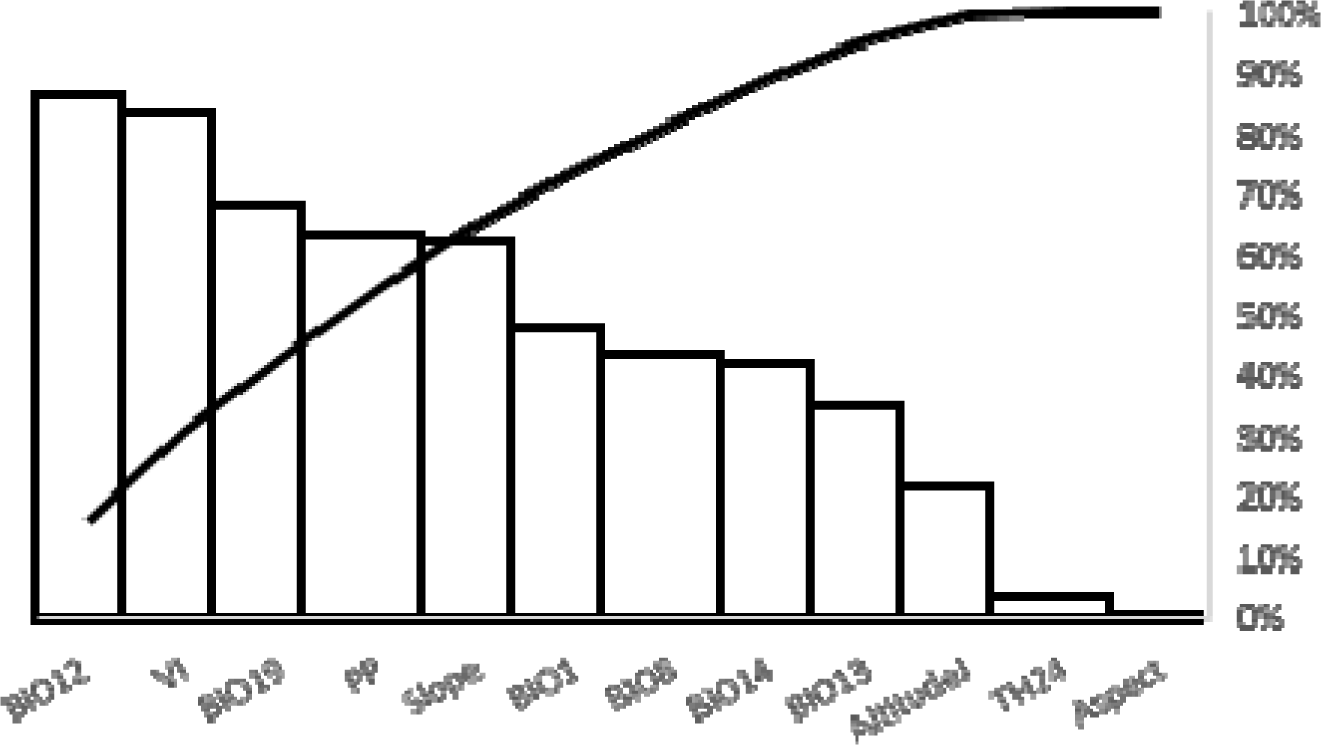
Variable contribution of *Diatraea* spp. distribution. BIO1 = Average Annual Temperature, BIO8 = Average Temperature of the Warmest Quarter, BIO10 = Average Temperature of the Warmest Quarter, BIO12 = Average Temperature of the Warmest Quarter, BIO12 = Annual Precipitation, BIO, VI = Normalized Vegetation Index; PP = Primay Production; BIO19 = Precipitation of Coldest Quarter; BIO13 = Precipitation of Wettest Month; BIO14 = Precipitation of Driest Month; TH24 = Topographical Heterogeneity

### Optimum niche

An optimal niche estimate where *Diatraea* spp. can express its maximum potential as a species, and indicates that under temperatures between 20°C and 23°C, accumulated annual rainfall between 1200 and 1500 mm, months where this variable is below 50 mm, slopes below 0.05, crop heterogeneity with an index of 0.2 and primary production values of 1.0, are ideal for settlement the ability of a crop to undergo compensatory growth can be influenced by the spatial pattern of injury. These conditions are similar to those present throughout the year in the Cauca river valley and the municipalities sampled in the department of Caldas; it is clear that the ranges obtained here have a greater or lesser amplitude for each environmental variable, facilitating the mobilization of pest populations towards different spaces than those currently occupied (Fig. 6).

**Fig 6.**
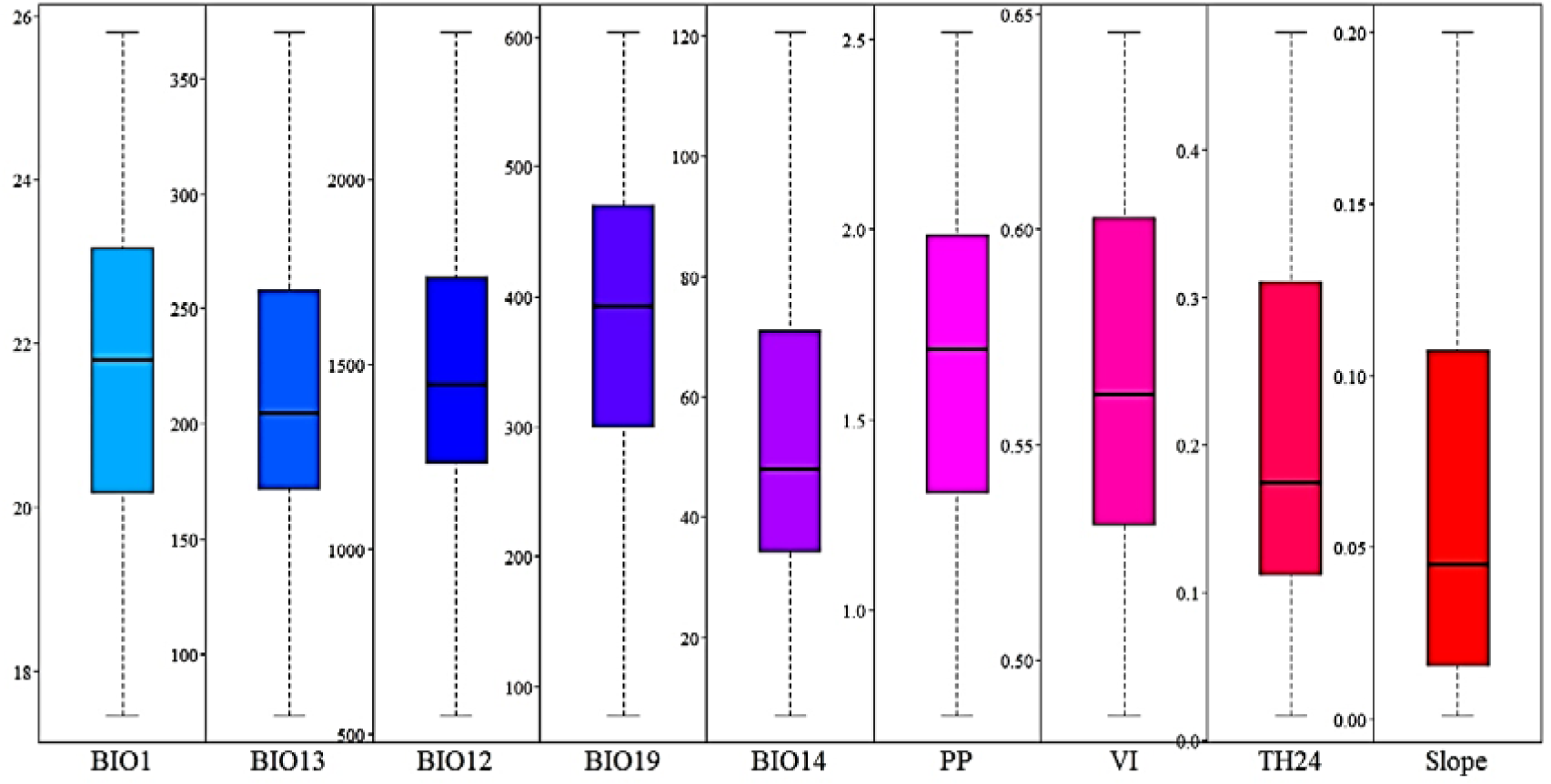
Estimation of an optimum niche for the establishment of *Diatraea* spp. BIO1 = Average Annual Temperature, BIO8 = Average Temperature of the Warmest Quarter, BIO10 = Average Temperature of the Warmest Quarter, BIO12 = Average Temperature of the Warmest Quarter, BIO12 = Annual Precipitation, BIO, VI = Normalized Vegetation Index; PP = Primay Production; BIO19 = Precipitation of Coldest Quarter; BIO13 = Precipitation of Wettest Month; BIO14 = Precipitation of Driest Month; TH24 = Topographical Heterogeneity

### Factors affecting *Diatraea* spp. richness

The most notable differences between one place and another have to do with the type of soil, the topography of the terrain, the altitude, the ambient temperature and the rainfall. These differences condition the distribution of flora and fauna. The matrix identifies the factors related to species richness, and their relative contribution, negative residues may be potential areas with lower species richness due to the negative effect of anthropogenic factors such as primary production (PP) and vegetation index (VI) (Fig. 7).

The polar coordinates determine the high probability of dispersion of *Diatraea* spp., where precipitation and slope are the determining environmental factors in the movement of adults to colonize new areas (Fig. 7).

## Distribution maps

The BCC_CSM1 climate change model, projected for 50 years, indicates a possible reduction in the number of niches and therefore in the population of *Diatraea* spp. under the premise of an increase in greenhouse gas concentrations and CO_2_, and while the temperature of the earth increases there will be an affection on the life cycle and dispersion due to meteorological alterations and changes in the ecosystem (Fig. 8).

**Fig. 1.**
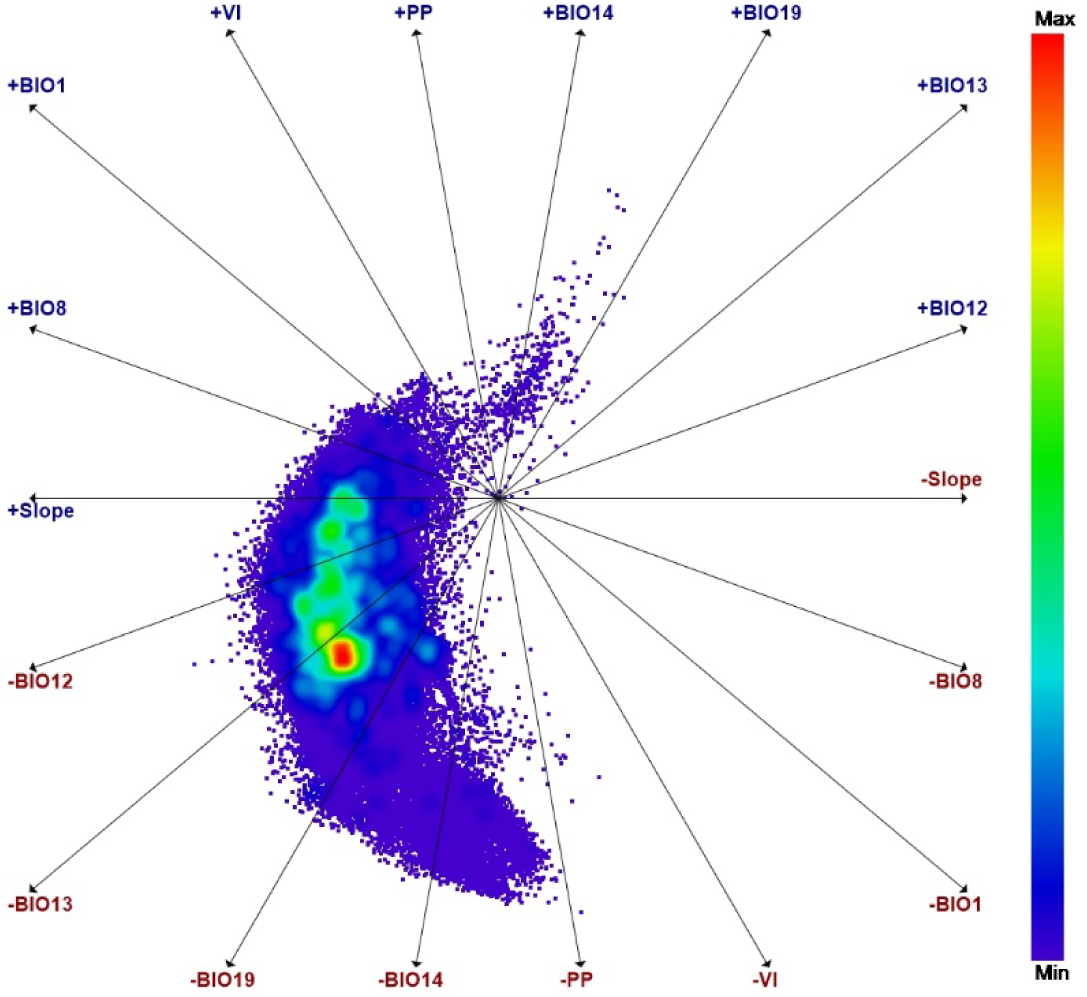
Polar coordinates to evaluate the importance of the environmental variable in the distribution of *Diatraea* spp. BIO1 = Average Annual Temperature, BIO8 = Average Temperature of the Warmest Quarter, BIO10 = Average Temperature of the Warmest Quarter, BIO12 = Average Temperature of the Warmest Quarter, BIO12 = Annual Precipitation, BIO, VI = Normalized Vegetation Index; PP = Primay Production; BIO19 = Precipitation of Coldest Quarter; BIO13 = Precipitation of Wettest Month; BIO14 = Precipitation of Driest Month; TH24 = Topographical Heterogeneity.

When climate change effects are projected over 70 years, the impact is clearly greater, with declines in *Diatraea* spp. populations in their current and potential niches of more than 50% when CO_2_ and greenhouse gas concentrations are projected to affect global temperature increases of up to 1°C, affecting not only the richness of plant species, water availability, but also development rates of insect populations, which depend on temperature as a whole (Fig 9).

**Fig. 2.**
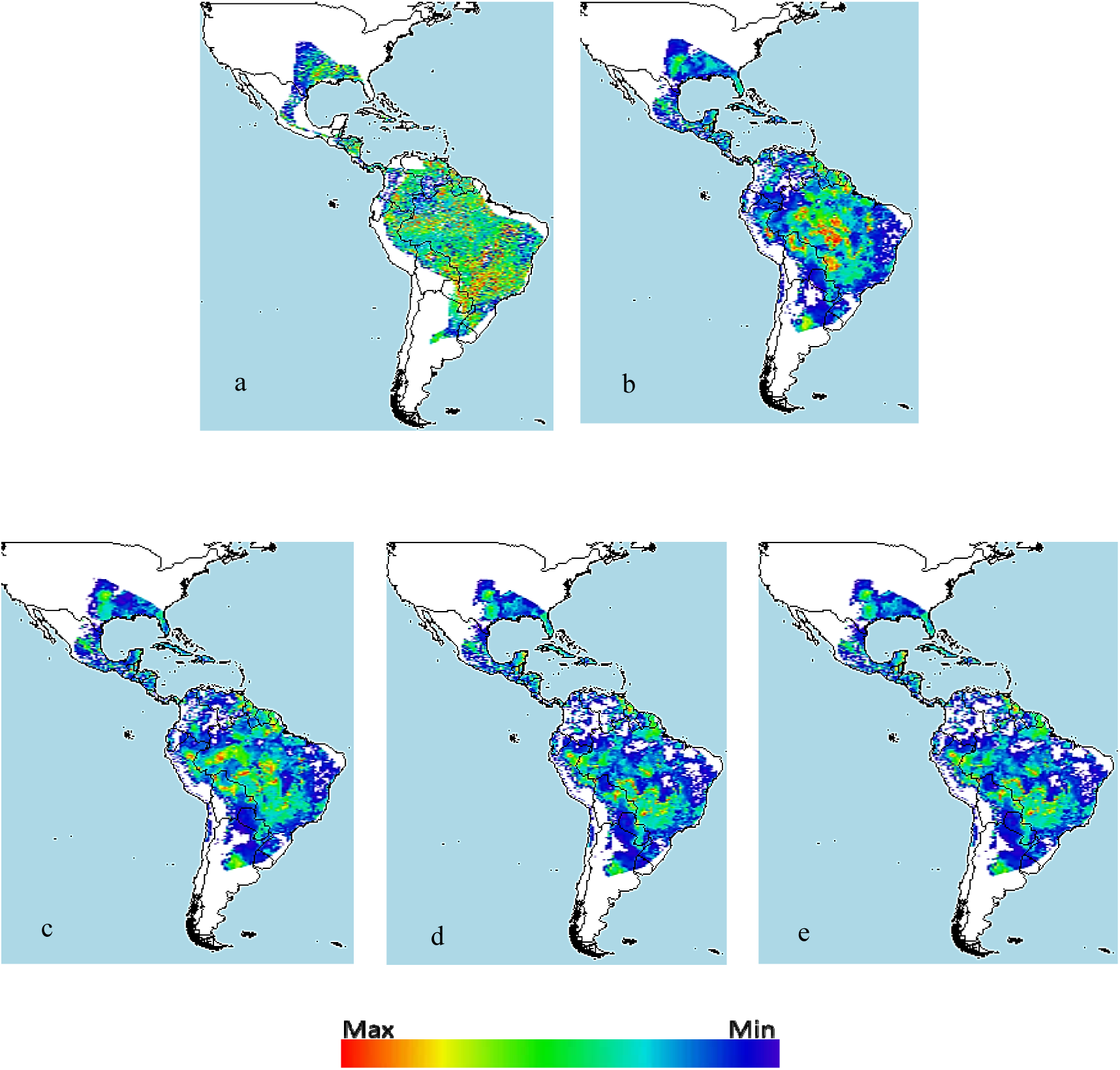
Distribution of *Diatraea* spp. in the western hemisphere under different climate change projections (BCC-CSM1-1) projected over 50 years at different carbon concentrations a) current condition, b) rcp26, c) rcp45, d) rcp60, e) rcp85. The color chart indicates maximum or minimum presence in a likely niche under favorable conditions.

**Fig. 3.**
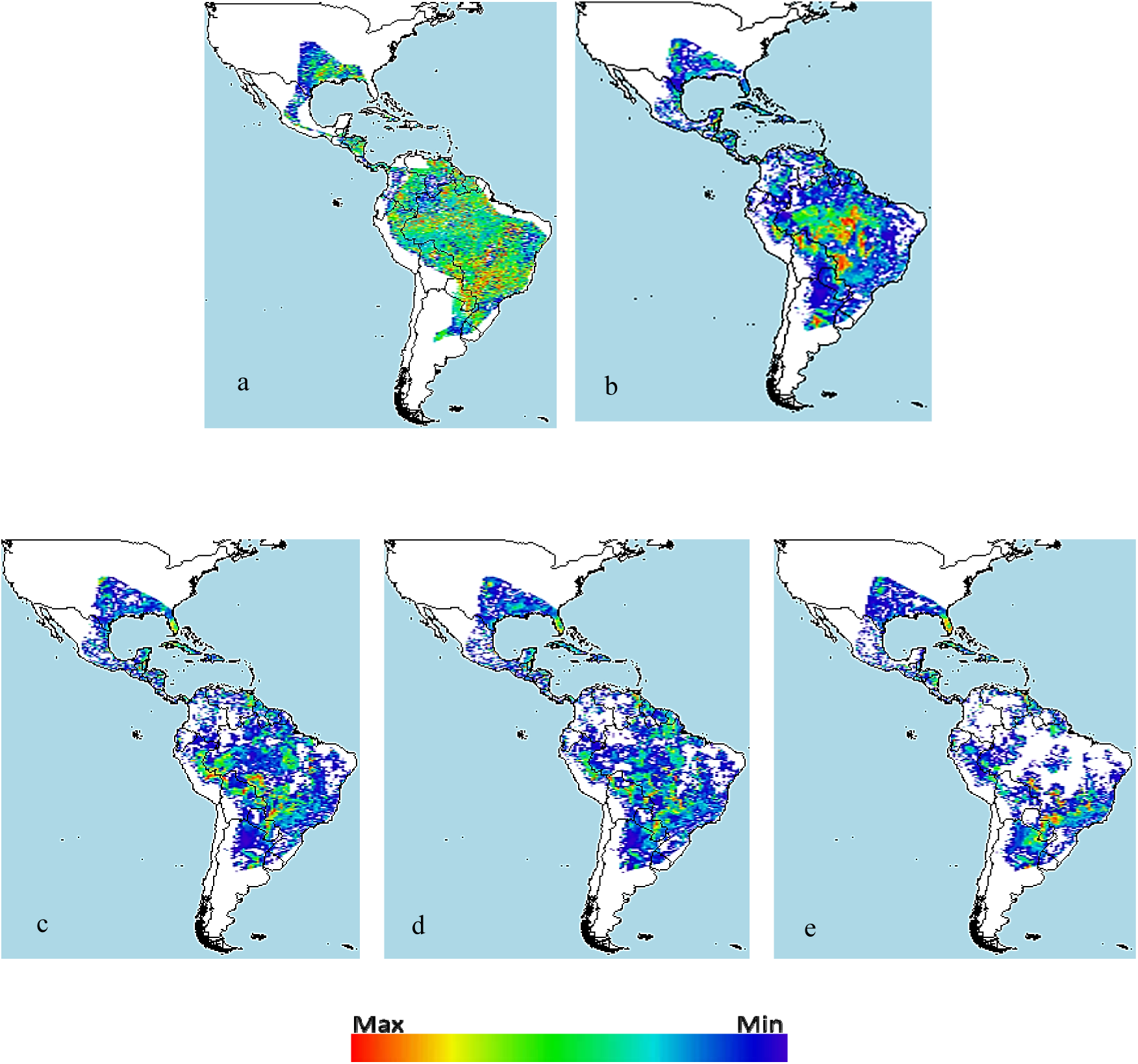
Distribution of *Diatraea* spp. in the western hemisphere under different climate change projections (BCC-CSM1-1) projected over 70 years at different carbon concentrations a) current condition, b) rcp26, c) rcp45, d) rcp60, e) rcp85. The color chart indicates maximum or minimum presence in a likely niche under favorable conditions.

At a more local scale, specifically the Colombian case, the direct effects of climate change on dispersion of *Diatraea* populations show population concentrations in some points such as the Caribbean and western regions of the country, completely changing the areas where they are currently found, specifically part of the mountain range of the western and central-southern mountain ranges of the country. The niche decrease may be greater than 50%, when projections under the climate change model are up to 50 years (Fig. 10). It should be noted that, although the *Diatraea* genus is a well-recognized sugarcane pest, it may be present in crops such as corn, rice, sorghum, and some forage grasses (Solis and Metz, 2016).

**Fig. 4.**
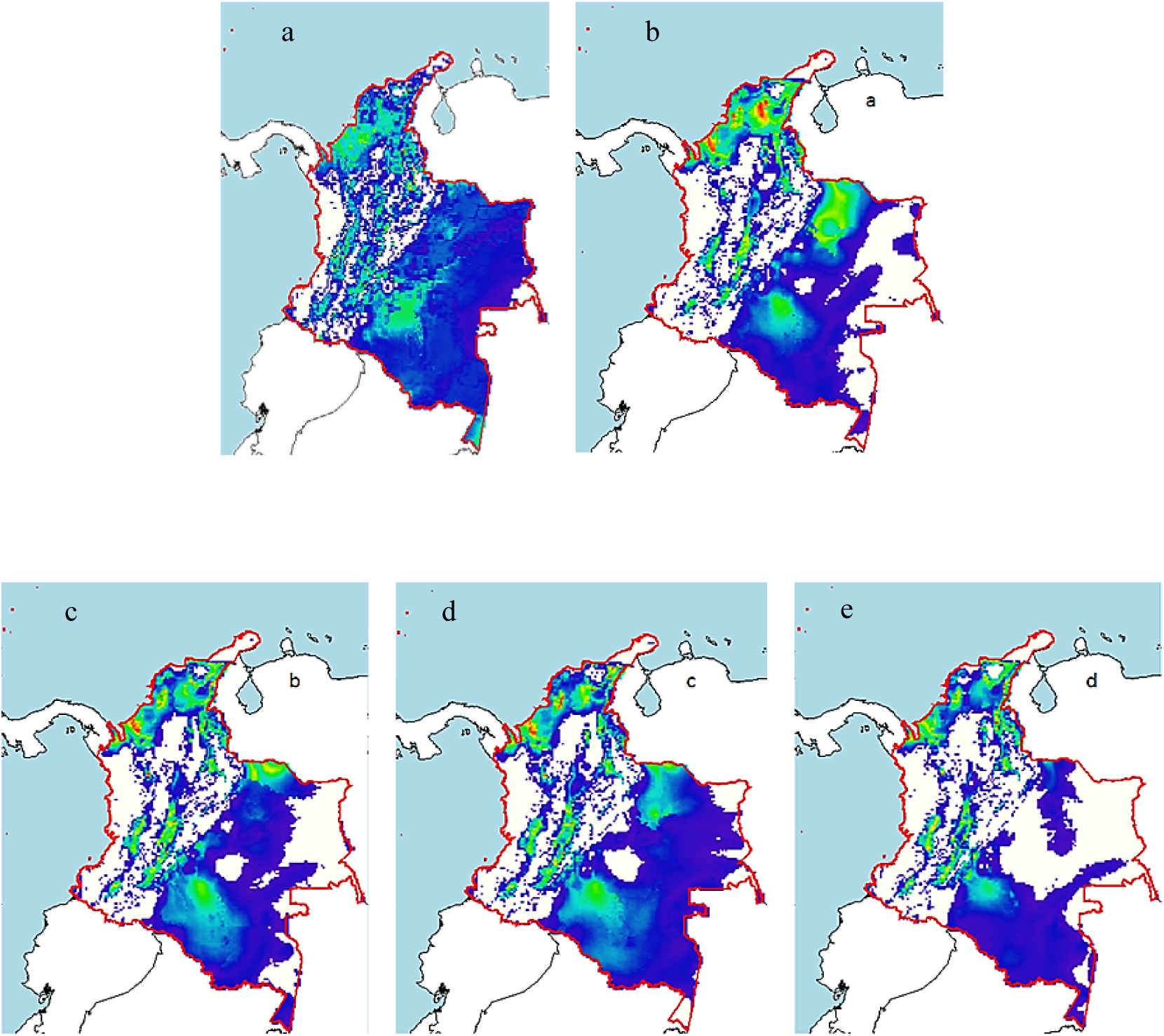
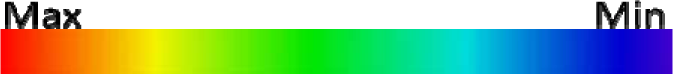
Distribution of *Diatraea* spp. in Colombia under different climate change projections (BCC-CSM1-1) projected over 50 years at different carbon concentrations a) current condition, b) rcp26, c) rcp45, d) rcp60, e) rcp85. The color chart indicates máximum or minimum presence in a likely niche under favorable conditions.

The models of maximum entropy and climate change can show reductions IN PEST POPULATIONS? of over 70%, where temperature increases can be the result of changes in water regimes and winds, radically changing climate dynamics at different scales (macro, meso and micro), focusing populations in smaller spaces, generating negative impacts on populations, and also on generation time rates (Fig. 11).

**Fig 5.**
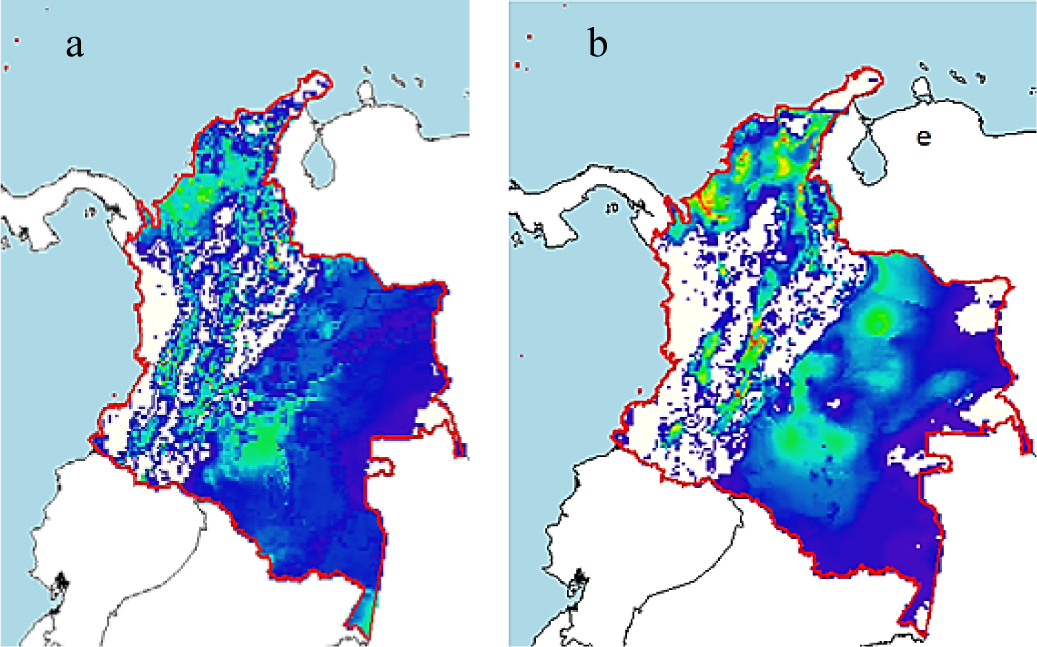
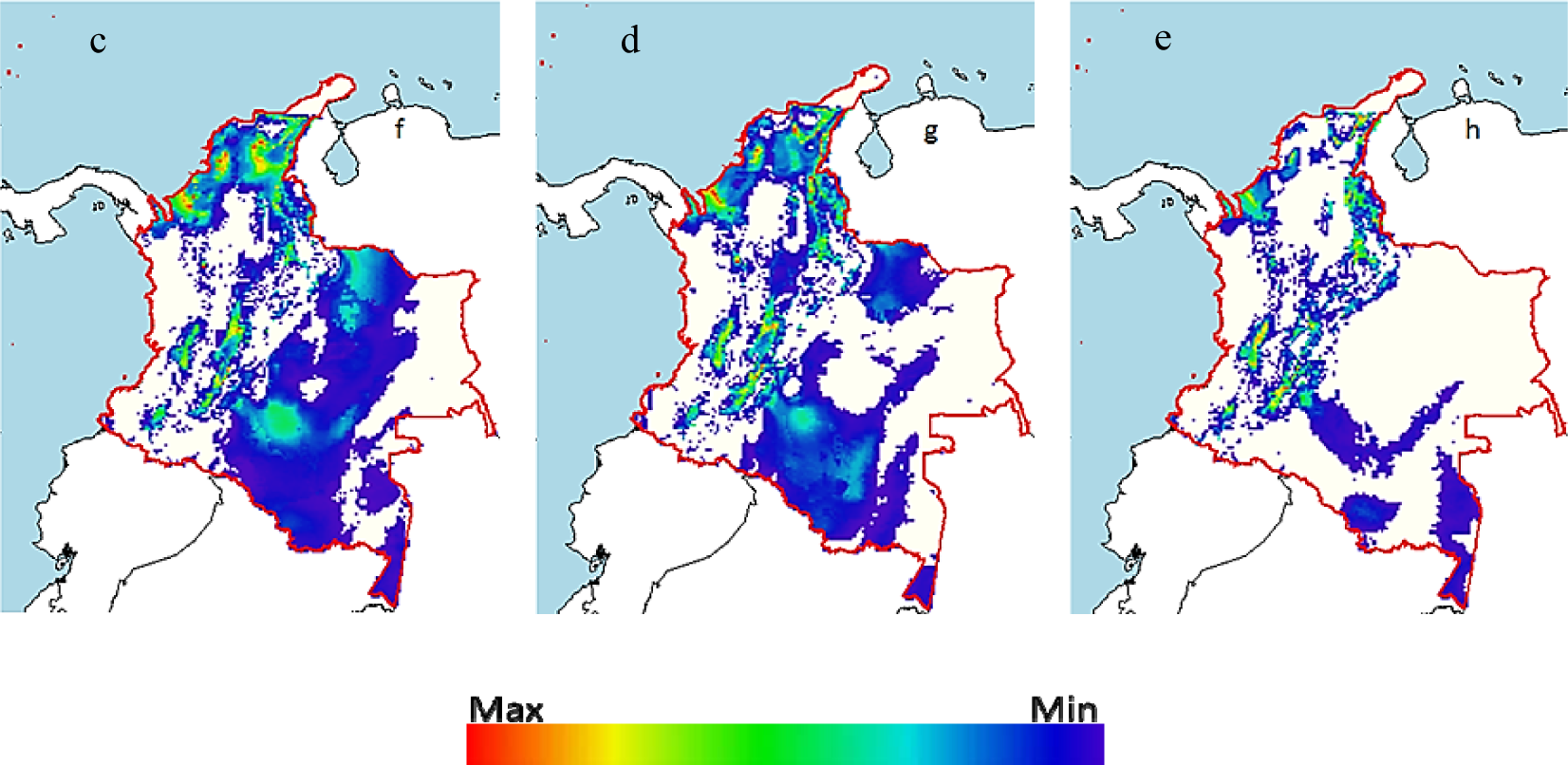
Distribution of *Diatraea* spp. in Colombia under different climate change projections (BCC-CSM1-1) projected over 70 years at different carbon concentrations a) current condition, b) rcp26, c) rcp45, d) rcp60, e) rcp85. The color chart indicates máximum or minimum presence in a likely niche under favorable conditions.

In relation to the Caldas department, it presents an important reduction not only of niche, but of populations, but they remained in some points of the central and western of Caldas, due perhaps to the positive effects that can bring the scenarios of heating that allow the permanence of the populations in this territory. The greatest effects on the development of *Diatraea busckella*. are evident throughout the Cauca river valley, where the highest concentration and production of sugarcane (*Saccharum officinarum*) is found, possibly due not only to atmospheric changes, but also to a possible need for change in the agricultural vocation of the territory (Fig 12).

**Fig 6.**
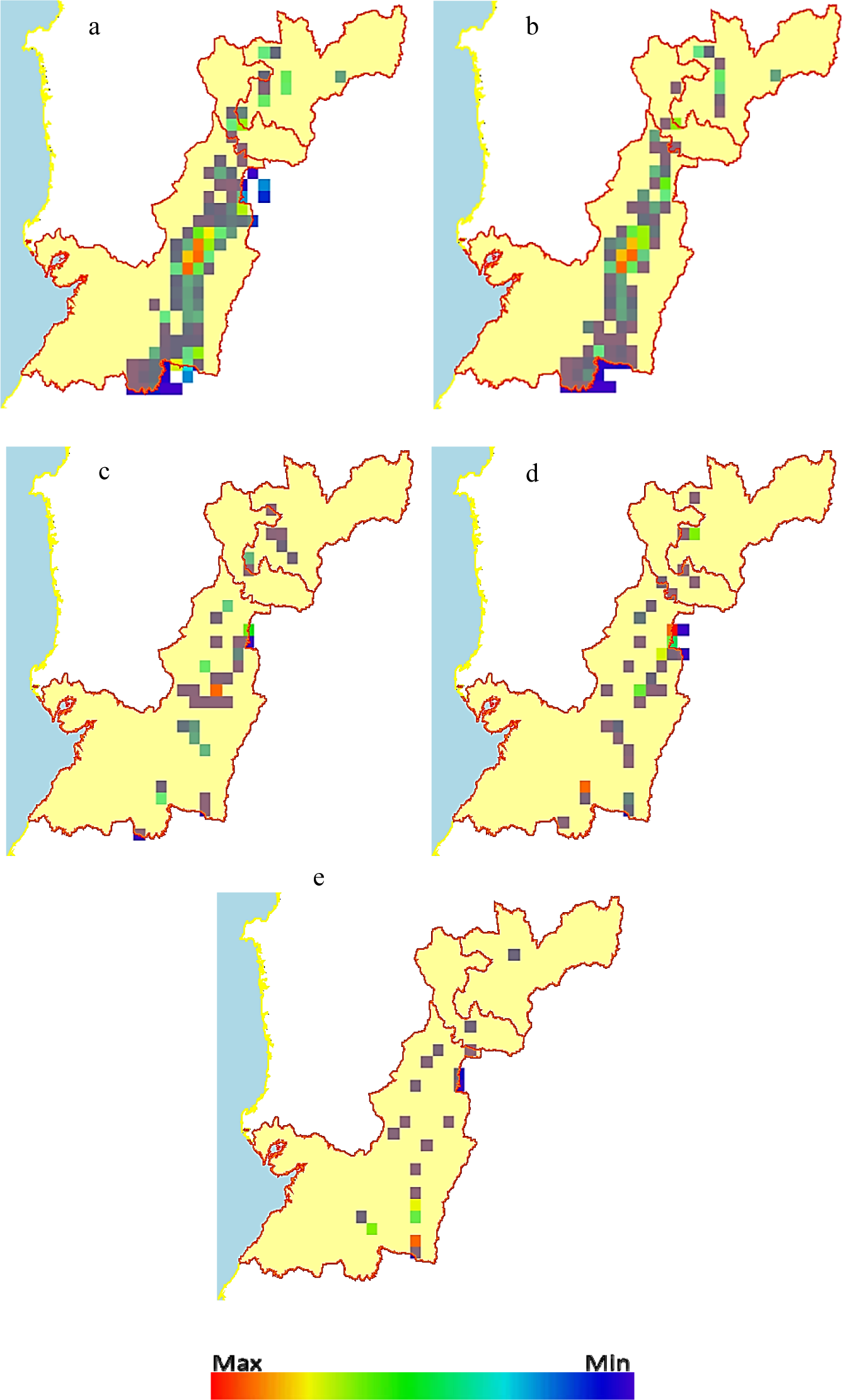
Distribution of *Diatraea busckella* under different climate change projections (BCC-CSM1-1) projected for 50 years in the departments of Caldas, Risaralda and Valle del Cauca a) current condition, b) rcp26, c) rcp45, d) rcp60, e) rcp85, in southwestern Colombia. The color chart indicates maximum or minimum presence in a likely niche under favorable conditions.

## Discussion

Climate data are widely used in the modeling of potential geographic distributions (Rojas *el al*., 2012, Mateo *el al*., 2013), as they allow the use of a greater number of historical records of species obtained in different years without incurring temporal inconsistencies between the records and the coverages used for modeling. In this way, it is intended to replace other types of coverings such as vegetation, which may limit the analyses since they require more updated records of the species that correspond only to the year in which they were developed (Plasencia *el al*., 2014).

The broadening of Geographic Information Systems and the growth of applied statistical techniques have allowed in recent years the expansion of tools for the analysis of spatial patterns of presence and absence of species(Franklin, 1995; Guisan & Zimmermann, 2000; Rushton *el al*., 2004; Foody, 2008; Swenson, 2008; Mateo *el al*., 2011) Species distribution models are necessary to evaluate the impacts not only on natural ecosystems, but also on agricultural ones, as well as to highlight the behavior of different insects species that are commonly associated to these areas, anticipating for those new colonizers, who have the potential for pest infestation.

The first models that want to describe the relationship between climatic phenomena, insect distribution and behavior were the linear relationships recorded by various authors (Candolle, 1885; Reibisch, 1902; Sanderson and Peairs, 1913; Arnold, 1960; Baskerville and Emin, 1969; Abrami, 1972; Allen, 1976, Sevacherian *et al*, 1977), which assumes the line as a valid relationship between development rate and temperature, to asymmetric (Janisch, 1925), Exponential (Belehradek, 1935) and Logistic (Davidson, 1944) models, to linear and maximum entropy models, in order to improve the calculation of the minimum development threshold. (Sharpe and DeMichele, 1977; Hilbert and Logan, 1983; Lactin *el al*., 1995; Briere *el al*., 1999; Phillips *et al*., 2006; Romo *et al*., 2013; Pérez and Liria, 2013). Among the evaluated methods are probabilistic models, used by Silva Carvalho (2011), in studies with *D. saccharalis*. For the pest frequency distribution study, where the data from each sample obtained in the previous phase will be adjusted by the Poisson distribution, which hypothesizes that all individuals have the same probability of occupying a place in any space and that the presence of one individual does not affect the presence of the other (Barbosa and Perecin, 1982), and the negative binomial distribution, where the occurrence of one individual limits the presence of neighboring individuals in the same place (Perecin and Barbosa, 1992).

Hulme *el al*., (2008); Roy *el al*., (2012) described the current exotic flora and fauna of Great Britain, reflecting a small subset of species resident in climatologically comparable regions for which important links have been established. Therefore, it is necessary to understand how climate change can influence future pathways for introduction of exotic species, where the natural dynamics in tropical areas, can be also affected by global warming due to alterations in the accumulation of day-grades for each physiological process. Colombia, although not influenced by seasons, but by patterns in the behavior of bimodal and unimodal rainfall in the Andean region, when El Niño or La Niña conditions are not present (Jaramillo *et al*., 2011), determines the distribution of insects, the area of dispersion and the possible establishment of climatic and environmentally optimal zones for the development of populations. Invertebrates are particularly sensitive to climatic conditions, and parameters such as temperature, precipitation, relative humidity and soil moisture have been shown to be useful in predicting important events in the growth of pest populations (Chen *et al*., 2014; Klapwijk *et al*., 2012).

Hulle *et al*., (2010) determined changes in pest profiles in South Australia; Hoffmann *et al*., (2008) concluded changes in the distribution of aphid flights across Europe; Nelson *et al*., (2013), highlighted the destabilization of the tea moth (*Adoxophyes honmai*), (Order: Lepidoptera; Family: Tortricidae) outbreak cycles in Japan, demonstrating the importance of studying environmental impacts on arthropod populations at a global scale. These organisms, due to their rapid development, are essential to highlight the changes in the changing climate supply; for example, *Diatraea* species can colonize new areas, specifically in the drier months, where precipitation is not a limiting factor for their distribution and where temperatures if ranging from 18 to 26°C, as thresholds for their development, can determine successful establishment as food availability is guaranteed (i.e. sugarcane).

The main expectation with respect to recent and projected global climate variability scenarios will influence the likelihood that species will be able to improve the use of available thermal energy for growth and reproduction. The latter represented in pests response to changes in temperature and expected rainfall. In this regard, Hulme *et al*., (2016) found increases in generalized fungal, plant and arthropod communities in Great Britain. Hill *et al*., (2012), concluded that distributions of the three *Penthaleus* species (Order: Trombidiformes; Family: Penthaleidae) in Australia are correlated with different climate variables, and adequate climate is likely to decrease in the future. Kocmankova *et al*., (2011), determined that the models suggest an expansion of the suitable habitat area for both pests in central Europe when assessing the distribution of *Leptinotarsa decemlineata* (Order: Coleoptera; Family: Chrysomelidae), a Colorado potato beetle and *Ostrinia nubilalis* (Order: Lepidoptera; Family: Crambidae), the European corn borer. Stoeckli *et al*., (2012) concludes that under future conditions of rising temperatures (2045-2074) in Switzerland, the risk of an additional generation of *Cydia pomonella* (Order: Lepidoptera; Family: Tortricidae), the apple moth will increase from 0-2% to 100% and there will be a two-week change in adult flights during the winter. Changes in phenology and voltinism will require a change in plant protection strategies. Gilbert *et al*. (2016) found that invasive populations of the African fig fly *Zaprionus indianus* (Order: Diptera; Family: Drosophilidae) in India show latitudinal clones indicative of rapid adaptation changes.

## Conclusions

In recent years, a new tool has become widespread that allows analysis of spatial patterns of the presence of organisms: species distribution models. These models are based on statistical and cartographic procedures which, based on real presence data, make it possible to infer potentially suitable areas according to their environmental characteristics. Data from natural history collections can be used for this purpose and thus acquire a new use.

Information about species distribution is increasingly important for a wide range of aspects of decision support and management, such as biodiversity studies, species protection, species reintroduction, alien species, prediction of potential effects of ecosystem loss or global climate change, etc. In fact, such information is crucial for the construction of species distribution models, as deficiencies in data quality lead to uncertainty in species distribution models. These models are important tools for understanding the relationship between species distribution and environmental parameters.

This study demonstrated that the ModestR software is a friendly, useful and valuable tool for users interested in the various areas of research and knowledge of biodiversity for its easy handling and accessibility (also free of charge), its efficiency in the management and generation of high quality information resolution and its ability to permanently update taxonomic information for the conduct of studies on the distribution of species present in all ecosystems of Colombia and the world.

*Diatraea* spp. is strongly influenced by the effects of climate change, not only on a global and regional scale, but also on a local scale, considerably reducing its population niches as well as the number of individuals, considering in the future (50 - 70 years) under the environmental models evaluated, reductions that may exceed 70% when compared to current records. The estimate of an optimal niche for the *Diatraea* species includes temperatures between 20°C and 23°C, accumulated annual rainfall between 1200 and 1500 mm, months with dry conditions, whose precipitation is below 50 mm, slopes below 0.05, crop heterogeneity with an index of 0.2 and primary production values of 1.0. These estimated conditions are similar to those present throughout the year in the Cauca river valley and the municipalities sampled in Caldas department.

The present research allows us to give a future approximation on the incidence of a complex arthropod complex that is a limiting factor for the cultivation of sugarcane, which may be present during global warming scenarios, reducing its populations, but remaining in currently inhabited spaces and colonizing new places, where conditions are optimal for development. The development of these models is intended to feed the support systems for decision making and to reduce the levels of uncertainty when looking for new management practices, control and establishment of new sugarcane projects in the country.

## Acknowledgements

To the Colombian Sugarcane Research Center, Cenicaña for its valuable contribution and collaboration in the supply, development and analysis of information.

To Research Center for Research, Innovation and Technology to the Sugarcane Sector of the Caldas Department BEKDAU, for contributing significantly to the development and execution of the project.

To Washington State University WSU, for guiding me in the development and analysis of each of the parameters evaluated

